# A most wanted list of conserved protein families with no known domains

**DOI:** 10.1101/207985

**Authors:** Stacia K. Wyman, Aram Avila-Herrera, Stephen Nayfach, Katherine S. Pollard

**Affiliations:** Gladstone Institutes, San Francisco, CA; University of California, Berkeley, CA; Lawrence Livermore National Laboratory, Livermore, CA; University of California, San Francisco, CA; DOE Joint Genome Institute, Walnut Creek, CA

## Abstract

The number and proportion of genes with no known function are growing rapidly. To quantify this phenomenon and provide criteria for prioritizing genes for functional characterization, we developed a bioinformatics pipeline that identifies robustly defined protein families with no annotated domains, ranks these with respect to phylogenetic breadth, and identifies them in metagenomics data. We applied this approach to 271 965 protein families from the SFams database and discovered many with no functional annotation, including >118 000 families lacking any known protein domain. From these, we prioritized 6 668 conserved protein families with at least three sequences from organisms in at least two distinct classes. These Function Unknown Families (FUnkFams) are present in Tara Oceans Expedition and Human Microbiome Project metagenomes, with distributions associated with sampling environment. Our findings highlight the extent of functional novelty in sequence databases and establish an approach for creating a “most wanted” list of genes to characterize.

---

Genome sequencing and metagenomics are producing unprecedented amounts of data. But elucidation of gene function has not kept pace with the volume of identified genes. Homology-based annotation methods predict domains and functions for many new protein coding and RNA genes. However, many sequenced genes do not have significant homology to experimentally characterized domains or gene families. To quantify this problem, we developed a bioinformatics approach to identify *bona fide* protein families with no annotation and then characterized these with respect to their phylogenetic range and abundance in metagenomes. The result is FUnkFams, a prioritized catalog of genes for experimental discovery of function.

Our pipeline begins with a database of gene families, filters out truncated sequences without a start and stop codon, assigns annotations to all sequences in each family using one or more annotation databases, and records the taxonomy of the organism from which each sequence derived (Sup Fig SI). In a second step, metagenomic sequencing reads from a user-defined collection of samples are mapped to protein families, resulting in an estimate of protein family abundance in each sample. These data are then used to organize and rank gene families based on their level of annotation, number of sequences, phylogenetic diversity, and distribution across metagenomes.

We applied this approach to discover the least annotated, most phylogenetically diverse full-length protein families in the SFams database (Sharpton et al., 2012) (Supplemental Text). We used SFams, because it was compiled in a comprehensive, automated fashion from thousands of diverse genome sequences, and we applied bioinformatics filters to remove small and truncated families. Specifically, we first identified 224 409 SFams with at least three unique, homologous, full-length protein sequences. We then annotated the sequences in these SFams using two curated and frequently updated sources of protein domains: the PFam database (Finn et al., 2014) and the NCBI Conserved Domain Database (CDD) (Marchler-Bauer et al., 2011). This analysis showed that the majority of protein families lack even a single domain annotated in PFam or CDD (N=118 607 SFams, 52.9% of total). These protein families without domain annotation are comprised of sequences from many branches of the cellular tree of life (Fig 1A). For further analysis and top prioritization we selected a subset of 6 668 protein families with no annotated domains and sequences from two or more taxonomic classes (Sup Fig 2A). We call these Function Unknown Families (FUnkFams)(Sup Table S1). Most FUnkFams (84.3%) are not in UniProt xref (UniProt, 2015), and those that are in UniProt are largely annotated as hypothetical or uncharacterized proteins (Sup Table S2).

**Figure 1.**
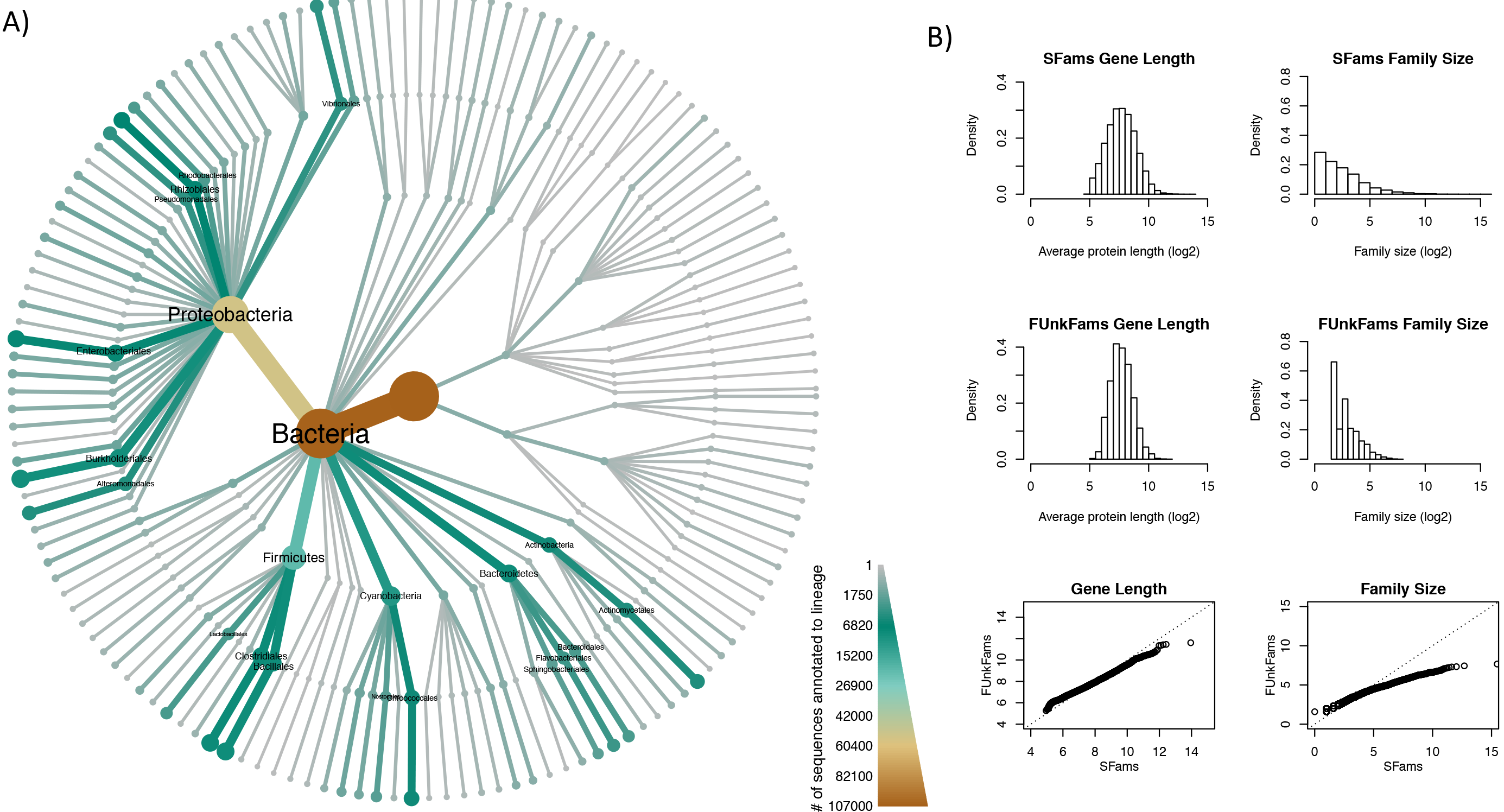
**A.** Phylogenetic heat tree of proteins in FUnkFams generated with Metacoder (Foster et al., 2017). Each FUnkFams protein sequence was annotated with the taxonomic label of the genome from which it was derived. The color of a branch represents the number of proteins from any FUnkFam on that branch of the taxonomy. The tree shows that FUnkFams are present across diverse lineages of cellular organisms including families from all three domains and over thirty phyla. Proteobacteria contribute many sequences to FUnkFams, in part because many genomes have been sequenced from that phylum. Supplemental Figure 2 shows the heat tree of all SFams, illustrating lineages where FUnkFams are enriched given how many genomes have been sequenced. **B.** FUnkFams protein length (in amino acids, log2 scale) and family size (number of protein sequences) are comparable to other SFams. Top and middle panels show histograms, and bottom panels are quantile-quantile plots showing that most quantiles of length and size are equal between FUnkFams and SFams, except at the top quan tiles where SFams are slightly longer (i.e., more amino acids) and bigger (i.e., more sequences).

FUnkFams are similar to other SFams in terms of properties other than the criteria we used to define them (i.e., functional annotation and phylogenetic breadth). Protein sequences in FUnkFams have a similar phylogenetic distribution to all SFams (Sup Fig S2B-C) with some enrichment in Cyanobacteria. They are also somewhat depleted in eukaryotes and archaea, probably due to bacterial SFams being more likely to meet our criteria of multiple homologous sequences from at least two classes. Like SFams, a typical FUnkFam is approximately 250 amino acids long (Fig 1B) and is comprised of three to five sequences (Fig 1C), though FUnkFams are slightly depleted for very long and very large families compared to better-annotated SFams. Nonetheless, six FUnkFams are comprised of more than 100 sequences, including a Proteobacterial family (SFams.ID=4560) with 203 sequences and a family (SFams.ID=5980) with 145 sequences spanning multiple domains of life. Thus, FUnkFams appear to be representative of full-length, phylogenetically diverse protein families.

To investigate the ecological distributions of FUnkFams, we quantified their abundance in shotgun metagenomes from the Tara Oceans Expedition (TO; 243 samples from 210 ecosystems in 20 biogeographic provinces at different depths over the course of three years) (Pesant et al., 2015) and Human Microbiome Project (HMP; 699 samples from oral, airways, skin, gut, vaginal sites on 300 healthy individuals at up to three time points over two years) Human Microbiome Project, 2012 (Supplemental Methods). The majority of FUnkFams (56.6%) are present in at least one of these 942 metagenomes, with many detected in multiple metagenomes (32.5% in at least two HMP samples, 37.2% in at least two TO samples) but relatively few (13.3%) detected in both TO and HMP (Fig 2A). FUnkFam prevalence was generally higher in TO (mean=18.6% versus 8.1%), with TO samples averaging 700 detected FUnkFams and HMP averaging 304 (Sup Fig S3A-B). Higher sequencing depth in TO may contribute to this signal. Abundance of detected FUnkFams is on average higher in TO, though many FUnkFams are approximately equally abundant between TO
and HMP (Fig 2B; Sup Fig S3C) and 27 are highly abundant in both environments (Sup Fig S4). Reflecting the ecological specificity of many FUnkFams, beta-diversity is significantly higher between the two environments than between samples within either environment (Fig 2C).

**Figure 2.**
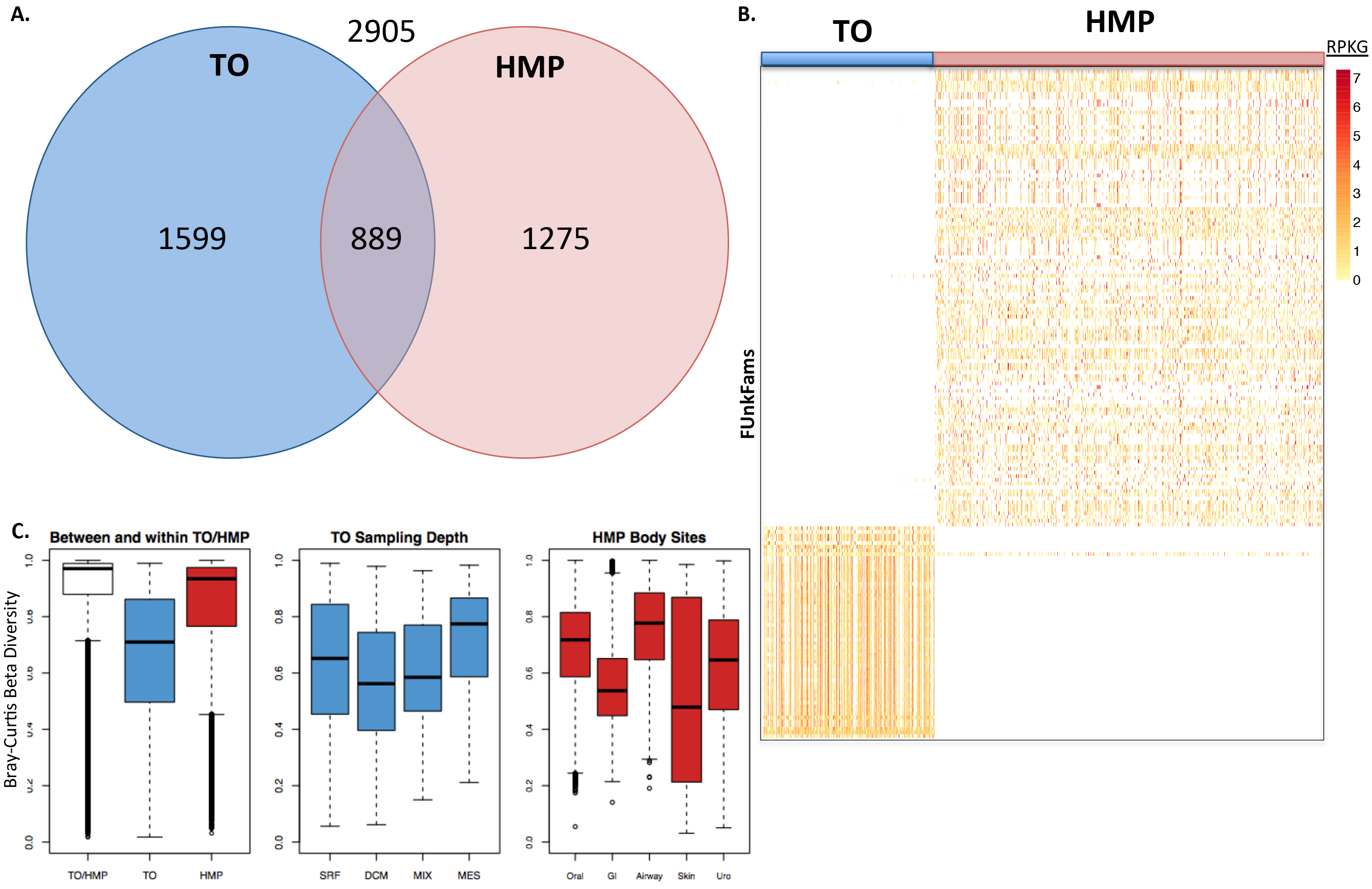
FUnkFams are present in marine and human metagenomes. **A.** Most FUnkFams are detected in either TO or HMP metagenomes, but relatively few are present in both environments. **B.** Heatmap showing the normalized abundance (RPKG) of FUnkFams (rows) in TO (left) or HMP (right) metagenomes. The 180 FUnkFams with at least 50 aligned reads across all samples are displayed (see Sup Fig S4 for the heatmap of all FUnkFams). **C.** Distributions of Bray-Curtis dissimilarity between pairs of samples from marine environments (TO; blue), between pairs of samples from human microbiomes (HMP; red), and between pairs of samples from different environments (white). Samples are more similar within than between the two environments.

We next used logistic regression to quantify how these differences in FUnkFam distributions across TO and HMP correlate with characteristics of the samples after adjusting for technical variables (Supplemental Methods). In TO, the presence of three FUnkFams was significantly associated with nitrate level after multiple testing correction (FDR<5%). One of these FUnkFams was also significantly associated with salinity and longitude, and another was significantly associated with longitude, latitude, temperature, and depth (Sup Table S3). Other FUnkFams showed weaker associations with environmental variables (Sup Fig S5, S6). The dominant variable associated with FUnkFam presence in HMP samples is body site (Sup Fig S5, S7; Sup Table S4), with only a few FUnkFams broadly detected across body sites. Other host phenotypes, such as BMI, smoking status, or diet, were not significantly associated with the presence of any FUnkFams.

These results identify thousands of uncharacterized protein families composed of homologous sequences from phylogenetically diverse organisms that are abundant in the human body or global oceans. These characteristics suggest that FUnkFams are *bona fide* protein families, and the associations of specific FUnkFams with marine environments or body sites provide hints about protein function and ecology. This study therefore lays the groundwork for significant future work to (i) predict (e.g., via genome proximity and further metagenome profiling (Lobb and Doxey, 2016) or literature based similarity (Price and Arkin, 2017)) and (ii) experimentally validate (e.g., via biochemical and structural characterization (Gerlt et al., 2011)) the functions of FUnkFams and the unannotated protein domains they contain. Our approach can be flexibly extended to use other databases of gene families and sources of functional annotation, and it will be interesting to apply it to other protein catalogs as well as RNA genes.

Supplementary information is available at ISME Journal’s website. FUnkFams data are freely available via figshare at: https://figshare.com/proiects/Function_Unknown_Families_of_homologous_proteins_FUnkFams_/25924.

## Acknowledgements

This project was funded by NSF (#DMS-1563159), Gordon & Betty Moore Foundation (#3300), and the Gladstone Institutes. We thank Thomas Sharpton and Jonathan Eisen for very helpful conversations at the conception of the project.

## Supplemental Text

### FUnkFams Construction

Our pipeline (Sup Fig S1) begins with 345 641 SFam protein families that were previously derived from *de novo* iterative clustering of ~10.5 million protein sequences from ~3 000 diverse genomes (Sharpton et al., 2012). SFams with less than three unique protein sequences are dropped, and then SFams where >50% of the sequences lack a start or stop codon are removed. This rigorous filtering produced 224 409 full-length protein families, at the cost of eliminating some small SFams. We then searched for all the proteins in these full-length SFams in the PFam database (acquired via Swissprot) (Finn et al., 2014) and the NCBI Conserved Domain Database (CDD) (Marchler-Bauer et al., 2011) to annotate domains in every sequence. These database searches were to identify the exact protein from an SFam (100% identical blast hit over the full length of the SFam sequence), not to identify homologs of the SFams sequences. The rationale for this strategy is that the SFam sequences derive from genomes that have been processed into these databases, and hence any proteins from these genomes should have been annotated already based on homology and other criteria of the databases. We identified 118 607 families for which no sequences had any domain annotation in PFam or CDD. Next, we characterized each sequence in each protein family according to the NCBI taxonomic annotation of the genome from which it derived and then quantified how many different species, genera, families, orders, classes, phyla, kingdoms, and domains are represented in each gene family (Sup Fig S2). We found that from the 118 607 families with no annotated domains there are 6 668 families that contain sequences from at least two classes. We call these FUnkFams. We identified 1 045 FUnkFams with at least one sequence in UniProt’s xref database (UniProt, 2015) (Sup Table S2). These are nearly all uncharacterized proteins, although we did identify eight FUnkFams with a sequence that has an xref-annotated function, despite having no domain annotation (Sup Table S5).

### Profiling in TO and HMP Metagenomes

We used Diamond (Buchfink et al., 2015) to align each read in the TO and HMP metagenome samples to a database of SFams sequences. We counted aligned reads for each FUnkFam, requiring a best hit to a protein belonging to the FUnkFam with at least 99% DNA sequence identity over the whole length of the read. FUnkFams with at least one read count were called present in the metagenome. FUnkFam abundance in each sample was estimated using reads per kilobase of genome (RPKG), a statistic that normalizes for both protein family length (mean of all member sequences in SFams database) and average genome size (estimated from the metagenomics sample with MicrobeCensus) (Nayfach and Pollard, 2015).

The TO dataset had hits to 2 488 FUnkFams in 308 samples, and the HMP dataset had hits to 2 164 FUnkFams in 696 samples (Fig 2A). Of these, 889 FUnkFams were detected in both environments, and these had a range of abundance levels across both TO and HMP (Sup Fig S3C). A particularly prevalent set of 137 FUnkFams was found in over 90% of TO samples, while just three were in over 90% of HMP samples, likely reflecting greater annotation of functions found in the human body samples relative to marine samples but also potentially also due to ecological differences between human body sites.

### Association Testing in HMP

We tested for association between a number of host phenotypes and FUnkFam presence in HMP metagenomes. To pre-filter FUnkFams without sufficient variation in presence across samples to detect associations, we only included FUnkFams with entropy in the top 25 percentile. To focus on the most phylogenetically diverse protein families, we additionally only included FUnkFams with sequences derived from genomes in at least two phyla (Sup Fig S2). This resulted in a set of 319 FUnkFams for association testing. We investigated associations with 13 host phenotypes that reflect lifestyle and medication use, as defined in HMP documentation (Sup Table S6). Phenotype data was obtained with permission through dbGaP (study ID = phs000228.v2.p1). Phenotypes were required to have at least two values with more than four observations. Seven subject variables passed this filtering step: bmi category, contraceptive use, breastfed status, diet, education level, birth country and student status. We fit a logistic regression model for each FUnkFam and used the resulting coefficients and their standard errors to perform t-tests to identify phenotypes associated with the presence of each FUnkFam across samples from each body site. The models account for geographic location (SITE variable in HMP) and were fit for each body site. P-values were corrected for multiple testing using the false discovery rate (FDR). We repeated this analysis within body subsites using the same filtering criteria and the resulting set of 335 FUnkFams (Sup Table S4).

### Association Testing in Tara Oceans Data

We tested for association between environmental variables and FUnkFam presence across TO samples. Environmental data was downloaded from the Tara Oceans data resource (http://ocean-microbiome.embl.de). Using the same criteria as with HMP, we analyzed only 100 FUnkFams with high entropy and sequences from at least two phyla. We fit logistic regression models for FUnkFam presence versus environmental variables, adjusting for latitude and month. Separate models were fit for samples collected with each filter size (size fraction). The resulting t-test p-values were adjusted for multiple testing using FDR (Sup Table S3).

## Supplemental Figures

**S1.** Bioinformatics pipeline for identifying FUnkFams from the SFams database

**S2.** A) Number of FUnkFams found across multiple domains, phyla, and classes in the tree of cellular organisms (e.g. 208 FUnkFams were found across more than one domain). B) Metacoder phylogenetic heat tree of SFams abundance across cellular organisms. Color indicates number of sequences on a branch. A random subset of 400 000 SFams was used to generate the tree. C) Metacoder phylogenetic heat tree of FUnkFams abundance across cellular organisms (as in Fig. 1A, for comparison here with SFams tree).

**S3.** A) Prevalence (vertical axis) of FUnkFams in TO (blue) and HMP (red) samples, ordered by decreasing prevalence in HMP (horizontal axis). B) Prevalence (vertical axis) of FUnkFams in TO (blue) and HMP (red) samples, ordered by decreasing prevalence in TO (horizontal axis). Many FUnkFams are more prevalent in TO than HMP, but the converse is not true. C) For 889 FUnkFams present in at least one TO and at least one HMP sample, the fractional abundance (vertical axis) represents the proportion of total RPKG for the FUnkFam that comes from TO (blue) versus HMP (red). FUnkFams are ordered by decreasing proportion of total RPKG deriving from TO samples (horizontal axis).

**S4.** Heatmap with all 3 763 FUnkFams (rows) detected in any metagenome (TO, HMP or both) at any abundance. Blue (left columns) are TO samples and red (right columns) are HMP samples.

**S5.** PCA plots of samples from HMP (A-B) and TO (C-E) based on counts of metagenomic sequencing reads mapped to all FUnkFams. HMP samples cluster by body site (A) but not other phenotypes such as BMI (B). TO samples cluster by marine layer (E) but not other environmental features (C-D).

**S6.** Heatmap for most abundant FUnkFams in TO samples, clustered both by column (samples) and row (FUnkFams) with environmental features annotated across rows.

**S7.** Heatmap for most abundant FUnkFams in HMP samples, clustered both by column (samples) and row (FUnkFams) with host phenotypes annotated across rows.

## Supplemental Tables

**Supplemental Table SI.** Characteristics of FUnkFams, including phylogenetic distribution and prevalence in TO and HMP samples.

**Supplemental Table S2.** Annotations for 1 045 FUnkFams with a protein sequence in the UniProt xref database.

**Supplemental Table S3.** Results of statistical tests for associations between environmental variables and FUnkFams presence across TO samples.

**Supplemental Table S4.** Results of statistical tests for associations between host phenotype variables and FUnkFams presence across HMP samples.

**Supplemental Table S5.** Annotations for eight FUnkFams with a protein sequence whose function is annotated in the UniProt xref database (despite having no annotated domains).

**Supplemental Table S6.** HMP phenotypes tested for association with FUnkFams abundance.

| HMP Variable | Description |
| --- | --- |
| DSUDIET | Meat/poultry frequency 294 |
| DSUBFED | Breastfed as child |
| BRTHCTRY | Born in USA or Canada |
| EDLVL_BS | Holds a BS degree or greater 295 |
| OCPTN_ST | Is a student |
| BMI_CAT | BMI categories |
| SMOKER | Uses tobacco and smokes cigarettes |
| DCMCODE_3 | Subject was taking antacids during the visit |
| DCMCODE_8 | Subject was taking antibiotics during the visit |
| DCMCODE_13 | Subject was taking contraceptives during the visit |
| DCMCODE_16 | Subject was taking GI meds during the visit |
| DCMCODE_18 | Subject was taking hormones/steroids during the visit |
| DCMCODE_NA | Subject was not taking medication during the visit |

